# Identification of consensus hairpin loop structure among the negative sense sub-genomic RNAs of SARS-CoV-2

**DOI:** 10.1101/2022.03.25.485826

**Authors:** Naveen Prakash Bokolia, Ravisekhar Gadepalli

## Abstract

SARS-CoV-2 is the causative agent of worldwide pandemic disease COVID-19. SARS-CoV-2 bears positive sense RNA genome, that have organized and complex pattern of replication/transcription process including the generation of subgenomic RNAs. Transcription regulatory sequences (TRS) have important role in the pausing of replication/transcription and generation of subgenomic RNAs. In the present bioinformatics analysis a consensus secondary structure was identified among negative sense subgenomic RNAs at the adjacent of initiation codon. This study proposed that consensus structured domain could involve in mediating the long pausing of replication/transcription complex and responsible for subgenomic RNA production.

## Introduction

COVID-19 is the worldwide pandemic disease caused by severe acute respiratory syndrome coronavirus 2 (SARS-CoV-2) that carries a positive sense, single strand RNA genome with the size of 29.9 kb (Kim et al., 2020; Zhu et al., 2020a). The RNA genome undergoes complex pattern of replication/transcription process inside the host environment. The SARS-CoV-2 enters the host cell and undergoes replication of positive sense RNA genome, where negative sense RNA genome is generated, that subsequently serve as template for positive sense RNA in order to translate the viral proteins and packaging of RNA genome in to virion particles. SARS-CoV-2 RNA genome bears conserved transcriptional regulatory sequences (TRS) of 6-7 nucleotides(Yang et al., 2020). The TRS is present at the immediate upstream of initiation codon, where replication/transcription complex is paused (Alexandersen et al., 2020; Kim et al., 2020). RdRp has the important property to backtracking that causes long pausing at specific site (Dulin et al., 2015; Malone et al., 2021). Therefore, in addition to complete RNA genome, a nested set of negative sense subgenomic RNAs is also generated including S subgenomic, 3a subgenomic, E subgenomic, M subgenomic, 6 subgenomic, 7a subgenomic, 7b subgenomic, 8 subgenomic, N subgenomic and 10 subgenomic. These canonical subgenomic RNAs are thought to be generated through the complex mechanism that involves pausing of negative sense RNA synthesis by RNA dependent RNA polymerase (RdRp) (Alexandersen et al., 2020; Kim et al., 2020; Mohammadi-Dehcheshmeh et al., 2021; Yang et al., 2020).

Although, the conserved role of TRS has been identified/proposed in the pausing of transcription, however, the role of RNA secondary structure has not been identified in this perspective. Therefore, in this study investigation of conserved secondary structure pattern was performed nearby the initiation codon/TRS that might be important in pausing of transcription during negative sense RNA synthesis of the SARS-CoV-2 genome.

The analysis of negative sense subgenomic RNAs was objectively performed in this study because of following important reason. The negative sense RNA is firstly generated through the replication/transcription of positive sense RNA genome. The negative sense RNA will serve as template for positive sense RNA genome and subgenomic messenger RNAs (or subgenomic RNAs) (Sawicki et al., 2007; Yang et al., 2020). According to previous research model, it was suggested that strand exchange or 5⍰ leader (5 ⍰ UTR) sequence to body (RNA) fusion occurs at TRS during negative sense RNA synthesis (Kim et al., 2020). Therefore, the consensus secondary structural motifs nearby the TRS might be important in several perspectives. In this study the bioinformatics analysis of negative sense subgenomic RNAs was performed that revealed a consensus hairpin loop secondary structure. Simultaneously, we validate the finding in parallel comparison with SARS-CoV and MERS-CoV. SARS-CoV-2 genome has 79.5% sequence similarity with SARS-CoV (Lu et al., 2020; Zhu et al., 2020b). Within this context it is significant to reveal the specific hairpin domain at the adjacent of initiation codon/TRS that is consistently present in SARS-CoV-2 and SARS-CoV subgenomic RNAs. In this perspective, research report proposed that consensus hairpin loop secondary structure during the synthesis of negative sense RNA mediates important role in transcriptional pausing and generation of subgenomic RNAs.

## Results

### The negative sense sub-genomic RNAs bear conserved hairpin domain at immediate downstream of initiation codon

Study by Alexandersen et al., 2021, determined the presence and abundance of subgenomic RNAs from the COVID-19 patient samples. Authors mapped the NGS reads to subgenomic RNAs in order to identify the abundance of particular subgenomic RNA from the patient samples(Alexandersen et al., 2020). NGS reads mapped results revealed that Orf7a and N subgenomic were in high abundance whereas Orf8, Orf6, and E subgenomic were relatively low (in an increasing order). These subgenomic RNAs are present several fold higher in comparison to whole genome fragments. Therefore, the present study seek to identify the possible conserved regions nearby the transcription regulatory sequences (TRS) or initiation codon those could mediate significant roles in the generation or expression of sub-genomic RNAs.

Therefore, in order to identify the consensus secondary structure, negative sense subgenomic RNAs were aligned by using LocARNA webserver. The significant length of upstream and downstream region (with respect to initiation codon) was considered for the analysis, with the total length of ∼420 nt. Sequence alignment revealed a consensus hairpin domain (∼35 to 60 nt; depending on the particular subgenomic RNA) among different negative sense sub-genomic RNAs (Figure 1, Table 1). This domain is present at the immediate downstream of initiation codon. The important features could be noted as the distance between initiation codon and consensus hairpin domain (Table 1). The structured hairpin domain of negative sense subgenomic RNAs might be of intrinsic functioning in either pausing /backtracking of RdRp at the immediate downstream of initiation codon. The conserved hairpin domain could also have role in template switching during transcription, although the role of conserved TRS has only been identified in recombination events (Yang et al., 2021). The conserved TRS is AACGAAC, that is highlighted in green (as GUUCGUU; negative sense) in the analyzed sequences of negative sense subgenomic RNAs (supplementary data file).

**Figure 1:**
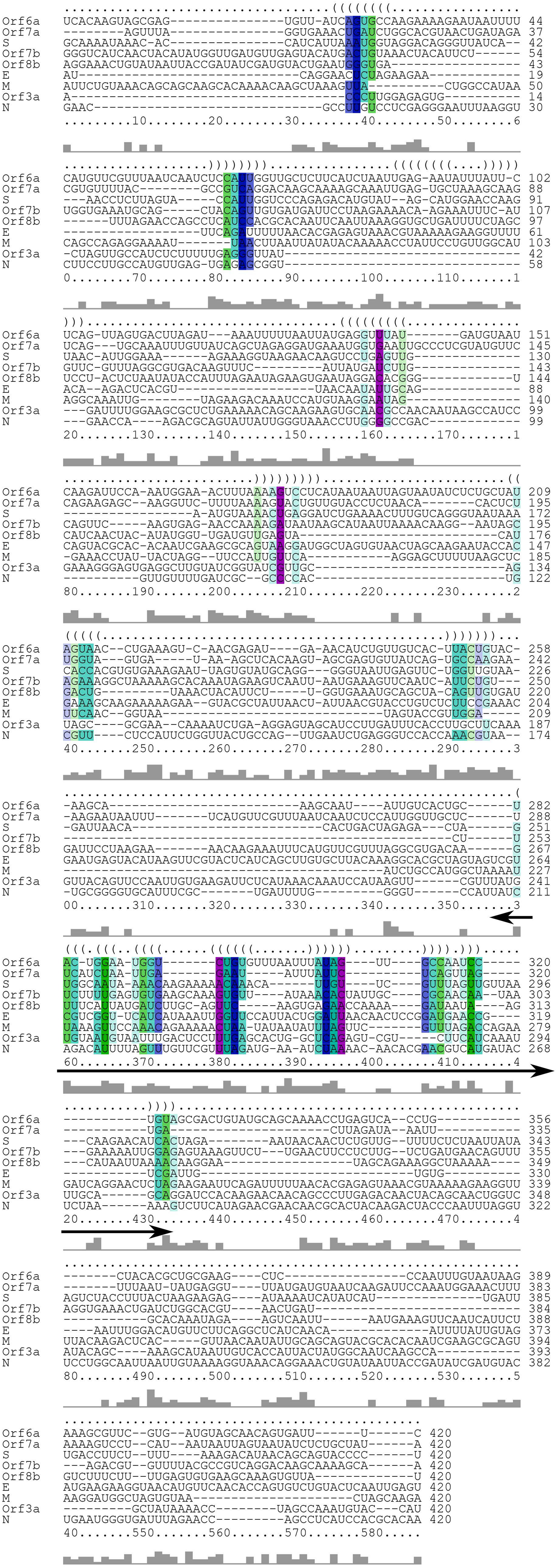
The figure presents the output of alignment result of negative sense subgenomic RNAs of SARS-CoV-2 that was done through the LocARNA webserver. The alignment result revealed the consensus sequence and hairpin structure (with double headed arrow) among negative sense subgenomic RNAs. This consensus structure is present at the immediate downstream of initiation codon or TRS (mentioned with double headed arrow).

**Table 1:** The table presents the description of analyzed negative sense subgenomic RNAs. The description is drawn from the combined output of sequence alignment followed by secondary structure prediction results. The number of asterisks (*) denotes to the abundance level for each subgenomic RNA detected by NGS reads mapped to the sequences and no asterisk is given to particular subgenomic that was detected at very low or near zero level) (Alexandersen et al., 2020).

| Sr. No. | Type of subgenomic RNA | Location of hairpin structure (from the downstream of initiation codon) | Length of hairpin loop structure with probability | Hairpin structure is distinct or not (merged with different loop carrying initiation codon) | Sequence alignment matches with predicted structure (probability) | Probability of formation of distinct secondary structure |
| --- | --- | --- | --- | --- | --- | --- |
| 1 | S subgenomic | Includes ~28 nt of coding region | ~55 nt | No (merged with initiation codon) | Yes | Less |
| 2 | *3a subgenomic | ~10 nt | ~58 nt | Yes | Yes | Moderate |
| 3 | ***E subgenomic | ~47 nt | ~52 nt | Yes | Yes | High |
| 4 | *M subgenomic | ~8 nt | Splits in three small hairpin loops | No (first loop merged with initiation codon) | No (three distinct hairpin structures) | Moderate |
| 5 | **Orf6 subgenomic | ~41 nt | ~47 nt | Yes | Yes | High |
| 6 | ****Orf7a subgenomic | ~30 nt | ~35 nt | Yes | Yes | High |
| 7 | Orf7b subgenomic | Includes ~6 nt of coding sequence | Splits in two hairpin loop structure | No (first loop merged with initiation codon) | No (two distinct hairpin structures) | Moderate |
| 8 | **Orf8 subgenomic | ~20 nt | ~57 nt | Yes | Yes | Moderate |
| 9 | ****N<br>subgenomic | ~1<sup>st</sup><br>nucleotide<br>of initiation<br>codon | ~60 nt | Yes | Yes | Moderate |

### The Orf7a subgenomic, N subgenomic, E subgenomic, Orf6 subgenomic and Orf8 subgenomic bears distinct secondary structure domain near the TRS

The sequence alignment of negative sense subgenomic RNAs revealed single hairpin domain. However, it was required to perform secondary structure prediction of each subgenomic RNA sequence to examine the possibility of consensus secondary structure formation. The secondary structure prediction was performed by using Vienna RNA webserver, and determined whether particular subgenomic RNA adopts a specific hairpin structure or not. Data obtained from sequence alignment and secondary structure prediction indicates that Orf7a subgenomic, N subgenomic, E subgenomic, Orf6 subgenomic and Orf8 subgenomic bears distinct and long hairpin domain with high probability of formation (Table 1, Figure 2A). These structural features could possibly be correlated with the previous study by Alexandersen et al; which described the high abundance of Orf7a subgenomic and N subgenomic RNAs whereas Orf8 subgenomic, Orf6 subgenomic, and E subgenomic were relatively low (Alexandersen et al., 2020). In addition, the remaining subgenomic RNAs (S subgenomic, Orf7b, Orf3a and M subgenomic) were reported to be very low or near zero level abundance (Alexandersen et al., 2020). The underlying reason of low level of NGS reads could be correlated with their respective hairpin structure (Figure 2B) and other properties mentioned in Table 1. These features include: hairpin loop splits in to subdomains, hairpin loop merged with initiation codon, length and probability of hairpin domain formation. Overall, the formation of distinct hairpin structure might be the underlying feature in the generation of negative sense sub-genomic RNAs.

**Figure 2:**
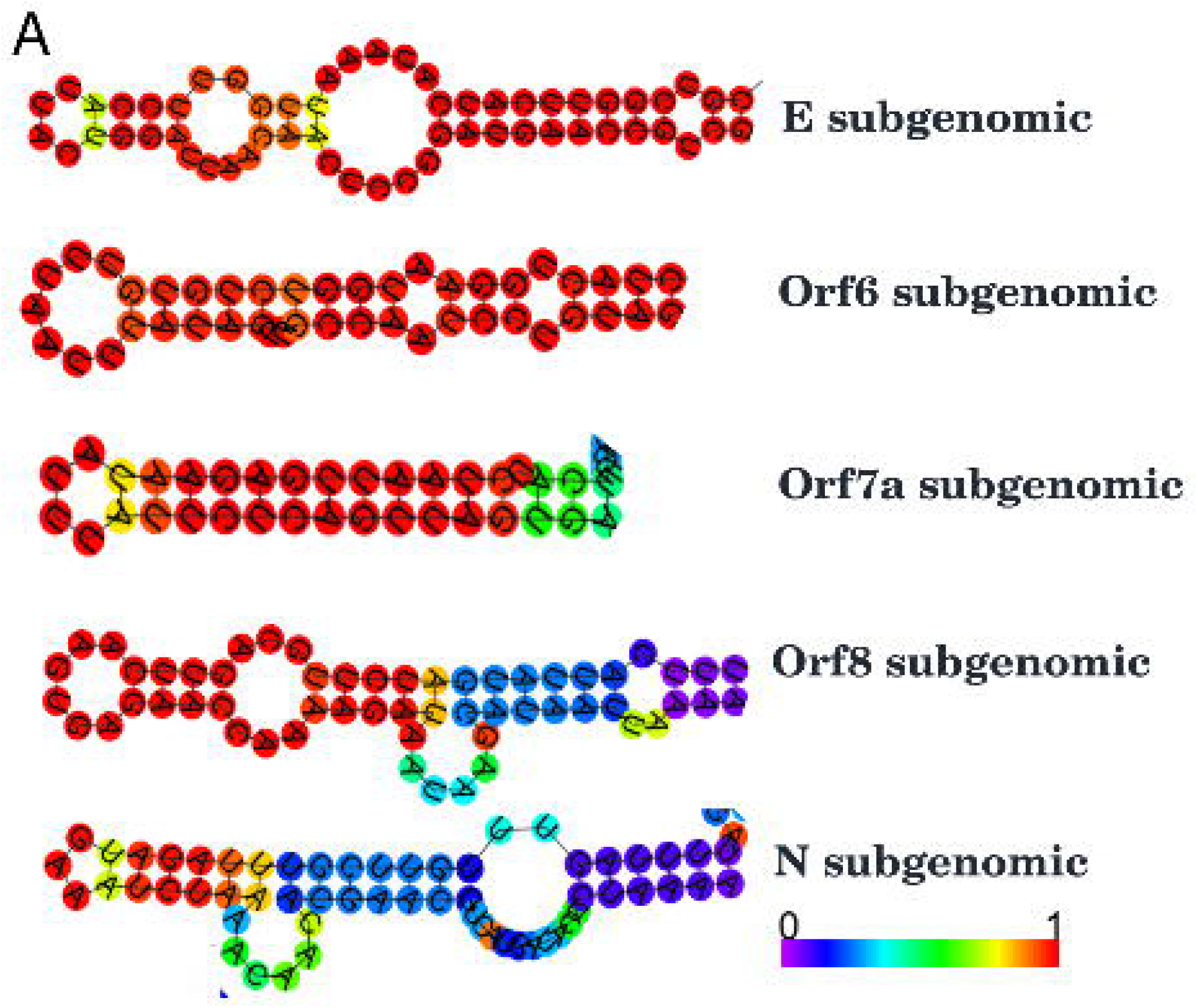
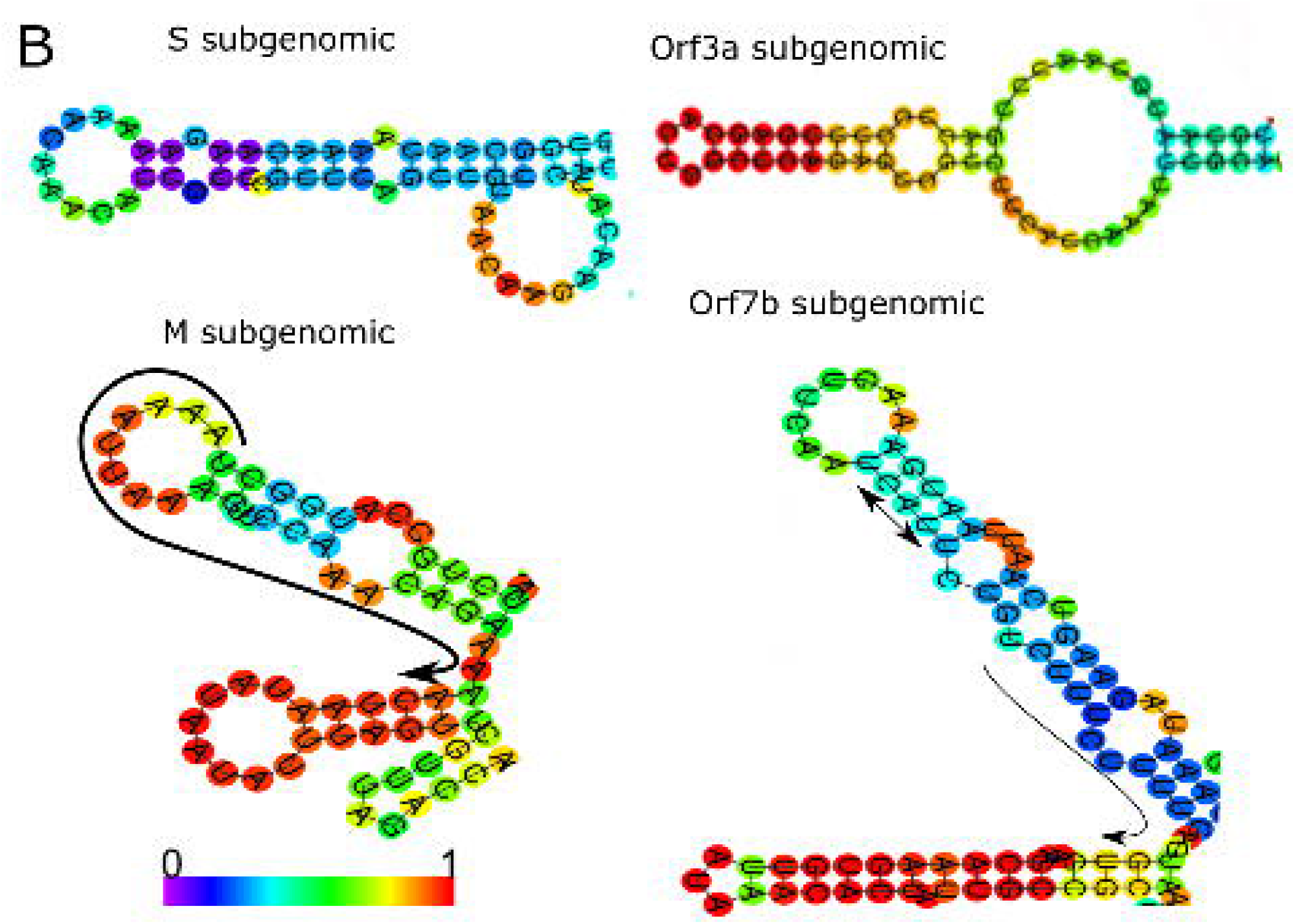
Secondary structure of consensus sequence of respective negative sense subgenomic RNAs. The secondary structure of each sequence was predicted by using Vienna RNA webserver. The respective portion of hairpin loop is shown with probability in color coding from 0 to 1. Secondary structure of consensus alignment sequence of each subgenomic RNA is shown in panel A and B. Panel A involves consensus secondary structure from subgenomic RNAs those have been detected or mapped at considerable level through NGS reads mapping (Alexandersen et al., 2020). Panel B involves consensus structure of subgenomic RNAs those have been detected at very low or zero level through NGS reads mapping. The complete secondary structure of each subgenomic RNA is provided in supplementary data file.

**Figure 3:**
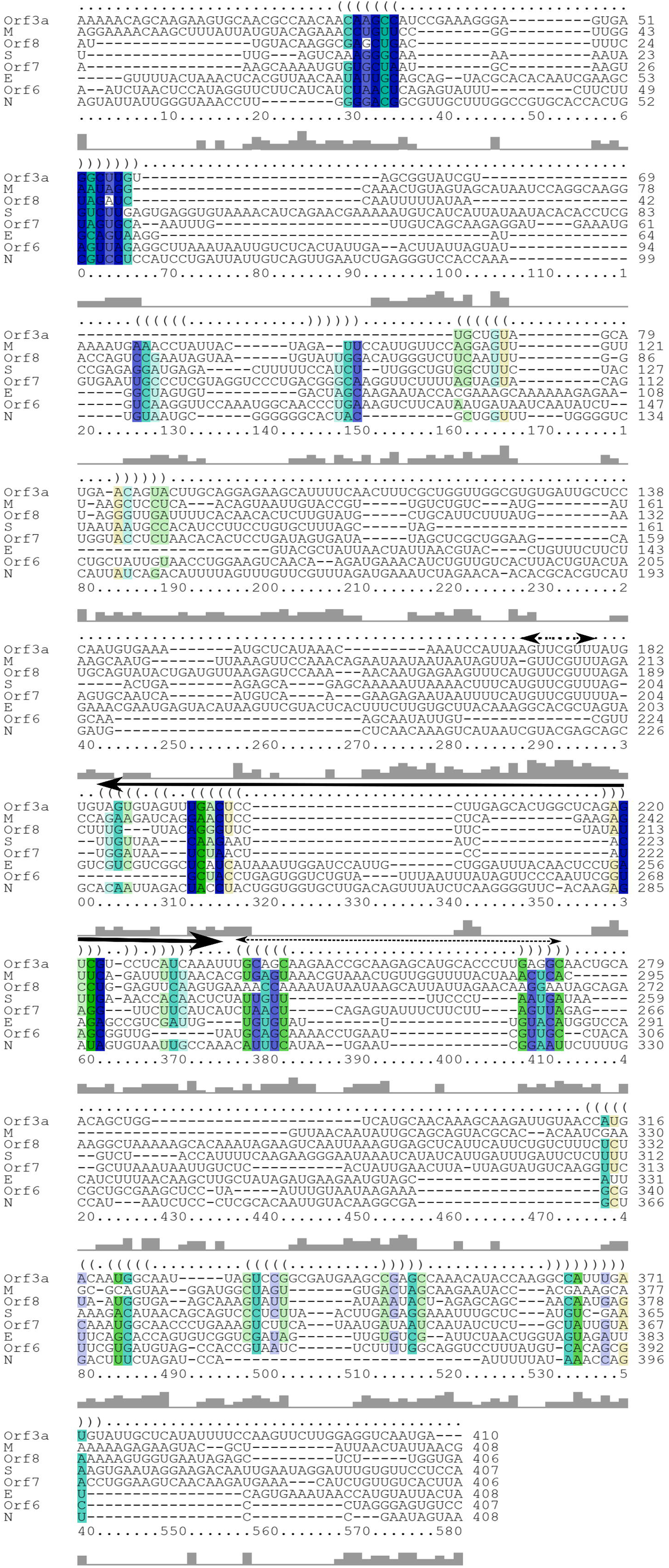
The figure presents the output of alignment result of negative sense subgenomic RNAs of SARS-CoV that was done through the LocARNA webserver. The alignment result revealed the consensus sequence and hairpin structure (with double headed arrow) among negative sense subgenomic RNAs. This consensus structure is present at the immediate downstream of initiation codon or TRS (mentioned with double headed arrow). In addition, the TRS sequence (GUUCGUU) is also shown just before the consensus hairpin domain.

Importantly, the highest similarity (and intermediate alignment) was observed between Orf7a and N subgenomic RNAs, as both subgenomic RNAs were placed together in guide tree. It provides significant relevance and correlation from the previously reported high abundant reads of N and Orf7a subgenomic (Alexandersen et al., 2020). In this perspective, we propose that conserved secondary structural elements might be a determining factor in transcriptional pausing and backtracking of RdRp that leads to subgenomic RNA production.

### Analysis of consensus secondary structure prediction in SARS-CoV subgenomic RNAs

It is important to validate or examine the possibility of consensus secondary structure in closely related enveloped viruses (beta-coronaviruses). Within this context, we further analyzed the subgenomic RNA sequences of SARS-CoV. We identified a distinct hairpin structure domain from the sequence alignment of subgenomic RNA sequences of SARS-CoV (Figure 4). The SARS-CoV subgenomic RNAs bears a distinct hairpin domain at the immediate downstream of TRS, as observed in the case of subgenomic RNAs of SARS-CoV-2 (Supplementary data file). However, after analysis it was observed that an additional domain could be considered in the case of some of the subgenomic RNAs of SARS-CoV as it falls at the immediate upstream of TRS, whereas for some of the specific subgenomic RNAs the second downstream domain could additionally be considered (as the consensus region falls at the immediate downstream of TRS).

### Discussion

The present finding revealed the conserved hairpin structure that is present in negative sense sub-genomic RNAs of SARS-CoV-2. The significance of conserved hairpin structure could be of two fold reasons. Firstly, the particular hairpin loop secondary structure could have intrinsic functioning in the pausing of replication/transcription of negative sense subgenomic RNAs mediated by RdRp. In addition, the conserved secondary structure could facilitate template switching (by unknown mechanism) at TRS during transcription, that involve the joining of 5⍰UTR to RNA body sequence. Further experimental studies could be performed to identify the much precise role of the conserved secondary structure nearby the initiation codon/TRS in the generation of negative sense subgenomic RNAs and maintenance of viral infection.

## Methods

### Extraction of SARS-CoV-2 RNA genome and subgenomic RNAs

The complete genome sequence of SARS-CoV-2 isolate Wuhan-Hu-1 (NCBI Reference Sequence: NC_045512.2) was used in this study, and specific regions (upstream and downstream) corresponding to particular subgenomic RNA were extracted. The nucleotides sequences were converted in to reverse complement that served as negative sense subgenomic RNA. The negative sense RNAs were used in alignment and secondary structure prediction analysis.

To analyze the subgenomic RNAs SARS-CoV, Bat coronavirus (BtCoV/279/2005), complete genome was used.

### RNA secondary structural similarity by using LocARNA webserver

LocARNA-Alignment and Folding webserver align the input RNA sequences and simultaneously fold them (Raden et al., 2018; Will et al., 2012). This webserver was used in the present study because it folds the RNA by using very realistic energy models as used by RNAfold of the Vienna RNA package. The sequences of negative sense subgenomic RNAs were aligned by using LocARNA webserver. The alignment type was global and in standard mode. The alignment was used to identify the consensus secondary structures elements between negative sense subgenomic RNA sequences of SARS-CoV-2.

### Secondary structure prediction by using Vienna RNA webserver

Secondary structure prediction of subgenomic RNA sequences was performed by using Vienna RNA webserver (Hofacker, 2003). The subset of aligned sequence region (∼285 nt) respective to each subgenomic RNA was submitted for structure prediction by using Vienna RNA webserver. The prediction results are included in supplementary data file.

