## Supplementary Data File 1 for "Identification of consensus hairpin loop structure among the negative sense sub-genomic RNAs of SARS-CoV-2"

**Negative sense Subgenomic RNAs sequences used in this study for alignment. Highlighted region: Initiation codon: yellow; TRS: Green; Consensus hairpin structure: Pink**

> S gene sub-genomic RNA
GCAAAAUAAACACCAUCAUUAAAUGGUAGGACAGGGUUAUCAAACCUCUUAGUACCAUUGGUCCCAGAG
ACAUGUAUAGCAUGGAACCAAGUAACAUUGGAAAAGAAAGGUAAGAACAAGUCCUGAGUUGAAUGUAA
AACUGAGGAUCUGAAAACUUUGUCAGGGUAAUAAACACCACGUGUGAAAGAAUUAGUGUAUGCAGGGG
GUAAUUGAGUUCUGGUUGUAAGAUUAACACACUGACUAGAGACUAGUGGCAAUAAAACAAGAAAAACA
AACAUUGUUCGUUUAGUUGUUAACAAGAACAUCACUAGAAAUAACAACUCUGUUGUUUCUCUAAUUA
UAAGUCUACCUUUACUAAGAAGAGAUAAAAUCAUAUCAUUGAUUUGACCUUCUUUAAAGACAUAACA
GCAGUACCCCU

> Orf3a Subgenomic
ACCCUUGGAGAGUGCUAGUUGCCAUCUCUUUUUGAGGGUUAUGAUUUUGGAAGCGCUCUGAAAAACA
GCAAGAAGUGCAACGCCAACAAUAAGCCAUCCGAAAGGGAGUGAGGCUUGUAUCGGUAUCGUUGCAGU
AGCGCGAACAAAAUCUGAAGGAGUAGCAUCCUUGAUUUCACCUUGCUUCAAAGUUACAGUUCCAAUUG
UGAAGAUUCUCAUAAACAAAUCCAUAAGUUCGUUUAUGUGUAAUGUAAUUUGACUCCUUUGAGCACUG
GCUCAGAGUCGUCUUCAUCAAAUUUGCAGCAGGAUCCACAAGAACAACAGCCCUUGAGACAACUACAGC
AACUGGUCAUACAGCAAAGCAUAAUUGUCACCAUUACUAUGGCAAUCAAGCCAGCUAUAAAACCUAGCC
AAAUGUACCAU

> E gene Subgenomic RNA
AUCAGGAACUCUAGAAGAAUUCAGAUUUUUAACACGAGAGUAAACGUAAAAAGAAGGUUUUACAAGAC
UCACGUUAACAAUAUUGCAGCAGUACGCACACAAUCGAAGCGCAGUAAGGAUGGCUAGUGUAACUAGCA
AGAAUACCACGAAAGCAAGAAAAAGAAGUACGCUAUUAACUAUUAACGUACCUGUCUCUUCCGAAACGA
AUGAGUACAUAAGUUCGUACUCAUCAGCUUGUGCUUACAAAGGCACGCUAGUAGUCGUCGUCGGUUCA
UCAUAAAUUGGUUCCAUUACUGGAUUAACAACUCCGGAUGAACCGUCGAUUGUGUGAAUUUGGACAUG
UUCUUCAGGCUCAUCAACAAUUUUAUUGUAGAUGAAGAAGGUAACAUGUUCAACACCAGUGUCUGUAC
UCAAUUGAGU

> M gene SUbgenomic RNA
AUUCUGUAAACAGCAGCAAGCACAAAACAAGCUAAAGUUACUGGCCAUAACAGCCAGAGGAAAAUUAAC
UUAAUUAUAUACAAAAACCUAUUCCUGUUGGCAUAGGCAAAUUGUAGAAGACAAAUCCAUGUAAGGAA
UAGGAAACCUAUUACUAGGUUCCAUUGUUCAAGGAGCUUUUUAAGCUCUUCAACGGUAAUAGUACCGU
UGGAAUCUGCCAUGGCUAAAAUUAAAGUUCCAAACAGAAAAACUAAUAUAAUAUUUAGUUCGUUUAGA
CCAGAAGAUCAGGAACUCUAGAAGAAUUCAGAUUUUUAACACGAGAGUAAACGUAAAAAGAAGGUUUU
ACAAGACUCACGUUAACAAUAUUGCAGCAGUACGCACACAAUCGAAGCGCAGUAAGGAUGGCUAGUGUA
ACUAGCAAGA

> Orf 6a Subgenomic
UCACAAGUAGCGAGUGUUAUCAGUGCCAAGAAAAGAAUAAUUUCAUGUUCGUUUAAUCAAUCUCCAU
UGGUUGCUCUUCAUCUAAUUGAGAAUAUUUAUUCUCAGUUAGUGACUUAGAUAAAUUUUUAAUUAUG
AGGUUUAUGAUGUAAUCAAGAUUCCAAAUGGAAACUUUAAAAGUCCUCAUAAUAAUUAGUAAUAUCUC
UGCUAUAGUAACCUGAAAGUCAACGAGAUGAAACAUCUGUUGUCACUUACUGUACAAGCAAAGCAAUA
UUGUCACUGCUACUGGAAUGGUCUGUGUUUAAUUUAUAGUUGCCAAUCCUGUAGCGACUGUAUGCAG
CAAAACCUGAGUCACCUGCUACACGCUGCGAAGCUCCCAAUUUGUAAUAAGAAAGCGUUCGUGAUGUAG
CAACAGUGAUUUC

> Orf 7a subgenomic RNA
AGUUUAGGUGAAACUGAUCUGGCACGUAACUGAUAGACGUGUUUUACGCCGUCAGGACAAGCAAAAGC
AAAUUGAGUGCUAAAGCAAGUCAGUGCAAAUUUGUUAUCAGCUAGAGGAUGAAAUGGUGAAUUGCCCU

CGUAUGUUCCAGAAGAGCAAGGUUCUUUUAAAAGUACUGUUGUACCUCUAACACACUCUUGGUAGUGA
UAAAGCUCACAAGUAGCGAGUGUUAUCAGUGCCAAGAAAAGAAUAAUUUUCAUGUUCGUUUAAUCAAU
CUCCAUUGGUUGCUCUUCAUCUAAUUGAGAAUAUUUAUUCUCAGUUAGUGACUUAGAUAAAUUUUUA
AUUAUGAGGUUUAUGAUGUAAUCAAGAUUCCAAAUGGAAACUUUAAAAGUCCUCAUAAUAAUUAGUAA
UAUCUCUGCUAUA

> Orf 7b Subgenomic RNA
GGGUCAUCAACUACAUAUGGUUGAUGUUGAGUACAUGACUGUAAACUACAUUCUUGGUGAAAUGCAGC
UACAGUUGUGAUGAUUCCUAAGAAAACAAGAAAUUUCAUGUUCGUUUAGGCGUGACAAGUUUCAUUA
UGAUCUUGCAGUUCAAGUGAGAACCAAAAGAUAAUAAGCAUAAUUAAAACAAGGAAUAGCAGAAAGGC
UAAAAAGCACAAAUAGAAGUCAAUUAAUGAAAGUUCAAUCAUUCUGUCUUUCUUUUGAGUGUGAAGCA
AAGUGUUAUAAACACUAUUGCCGCAACAAUAAGAAAAAUUGGAGAGUAAAGUUCUUGAACUUCCUCUU
GUCUGAUGAACAGUUUAGGUGAAACUGAUCUGGCACGUAACUGAUAGACGUGUUUUACGCCGUCAGGA
CAAGCAAAAGCAA

> Orf 8b SUbgenomic RNA
AGGAAACUGUAUAAUUACCGAUAUCGAUGUACUGAAUGGGUGAUUUAGAACCAGCCUCAUCCACGCAC
AAUUCAAUUAAAGGUGCUGAUUUUCUAGCUCCUACUCUAAUAUACCAUUUAGAAUAGAAGUGAAUAGG
ACACGGGUCAUCAACUACAUAUGGUUGAUGUUGAGUACAUGACUGUAAACUACAUUCUUGGUGAAAUG
CAGCUACAGUUGUGAUGAUUCCUAAGAAAACAAGAAAUUUCAUGUUCGUUUAGGCGUGACAAGUUUCA
UUAUGAUCUUGCAGUUCAAGUGAGAACCAAAAGAUAAUAAGCAUAAUUAAAACAAGGAAUAGCAGAAA
GGCUAAAAAGCACAAAUAGAAGUCAAUUAAUGAAAGUUCAAUCAUUCUGUCUUUCUUUUGAGUGUGAA
GCAAAGUGUUAU

> N gene subgenomic RNA
GAACGCCUUGUCCUCGAGGGAAUUUAAGGUCUUCCUUGCCAUGUUGAGUGAGAGCGGUGAACCAAGAC
GCAGUAUUAUUGGGUAAACCUUGGGGCCGACGUUGUUUUGAUCGCGCCCCACUGCGUUCUCCAUUCUG
GUUACUGCCAGUUGAAUCUGAGGGUCCACCAAACGUAAUGCGGGGUGCAUUUCGCUGAUUUUGGGGU
CCAUUAUCAGACAUUUUAGUUUGUUCGUUUAGAUGAAAUCUAAAACAACACGAACGUCAUGAUACUCU
AAAAAGUCUUCAUAGAACGAACAACGCACUACAAGACUACCCAAUUUAGGUUCCUGGCAAUUAAUUGUA
AAAGGUAAACAGGAAACUGUAUAAUUACCGAUAUCGAUGUACUGAAUGGGUGAUUUAGAACCAGCCUC
AUCCACGCACAA

### SARS CoV Negative Sense subgenomic RNAs with consensus region

>S subgenomic

UUUGAGUCAAAGGGCAAAAAAUAGUCUUGAGUGAGGUGUAAAACAUCAGAACGAAAAAUGUCAUCAUU
AUAAUACACACCUCGCCGAGAGGAUGAGACUUUUUCCAUCUUUGGCUGUGGCUUUCUACUAAUAAUGC
CACAUCCUUCCUGUGCUUUAGCUAGACUGAAGAGCAGAGCAAAAAUUAAAACUUUCAUGUUCGUUUAG
UUGUUAACAAGAAUAUCACUUGAAACCACAACUCUAUUGUUUUCCCUAAUGAUAAGUCUACCAUUUUC
AAGAAGGGAAUAAAUCAUAUCAUUGAUUUGAUUCUCUUUUAAAGACAUAACAGCAGUCCCUCUUAACU
UGAGAGGAAAUUUGCUCAUGUCGAAAAGUGAAUAGGAAGACAAUUGAAUAGGAUUUGUGUUCCUCCA

>Orf3a Subgenomic

AAAAACAGCAAGAAGUGCAACGCCAACAACAAGCCAUCCGAAAGGGAGUGAGGCUUGUAGCGGUAUCGU
UGCUGUAGCAUGAACAGUACUUGCAGGAGAAGCAUUUUCAACUUUCGCUGGUUGGCGUGUGAUUGCU
CCCAAUGUGAAAAUGCUCAUAAACAAAUCCAUUAAGUUCGUUUAUGUGUAGUGUAGUUUGACUCCCUU
GAGCACUGGCUCAGAGUCGUCCUCAUCAAAUUUGCAGCAAGAACCGCAAGAGCAUGCACCCUUGAGGCA

ACUGCAACAGCUGGUCAUGCAACAAAGCAAGAUUGUAACCAUGACAAUGGCAAUUAGUCCGGCGAUGAA
GCCGAGCCAAACAUACCAAGGCCAUUUGAUGUAUUGCUCAUAUUUUCCAAGUUCUUGGAGGUCAAUGA

>E subgenomic

GUUUUACUAAACUCACGUUAACAAUAUUGCAGCAGUACGCACACAAUCGAAGCGCAGUAAGGAUGGCUA
GUGUGACUAGCAAGAAUACCACGAAAGCAAAAAAGAGAAGUACGCUAUUAACUAUUAACGUACCUGUU
UCUUCUGAAACGAAUGAGUACAUAAGUUCGUACUCACUUUCUUGUGCUUACAAAGGCACGCUAGUAGU
CGUCGUCGGCUCAUCAUAAAUUGGAUCCAUUGCUGGAUUUACAACUCCUGAAGAGCCGUCGAUUGUGU
GUAUUUGUACAUGGUCCACAUCUUUAACAAGCUUGCUAUAGAUGAAGAAUGUAGCAUUUUCAGCACCA
GUGUCGGUCGAUAGUUGUGUCGAUUCUAACUGGUAGUAGAUUUCAGUGAAAUAACCAUGUAUUACUA

>M Subgenomic

AGGAAAACAAGCUUUAUUAUGUACAGAAACCUGUUCCGGUUGGAAUAGGCAAACUGUAGUAGCAUAAU
CCAGGCAAGGAAAAUGAAACCUAUUACUAGAUUCCAUUGUUCCAGGAGUUGUUUAAGCUCCUCAACAG
UAAUUGUACCGUUGUCUGUCAUGAUAAGCAAUGUUAAAGUUCCAAACAGAAUAAUAAUAAUAGUUAGU
UCGUUUAGACCAGAAGAUCAGGAACUCCCUCAGAAGAGUUCAGAUUUUUAACACGUGAGUAAACGUAA
ACUGUUGGUUUUACUAAACUCACGUUAACAAUAUUGCAGCAGUACGCACACAAUCGAAGCGCAGUAAG
GAUGGCUAGUGUGACUAGCAAGAAUACCACGAAAGCAAAAAAGAGAAGUACGCUAUUAACUAUUAACG

>Orf6 subgenomic

AAUCUAACUCCAUAGGUUCUUCAUCAUCUAACUCAGAGUAUUUCUUCUUAGUUAGAGGCUUAAAUAAU
UGUCUCACUAUUGAACUUAUUAGUAUGUCAAGGUUCCAAAUGGCAACCCUGAAAGUCUUCAUAAUGAU
AAUCAAUAUCUCUGCUAUUGUAACCUGGAAGUCAACAAGAUGAAACAUCUGUUGUCACUUACUGUACU
AGCAAAGCAAUAUUGUCGUUGCUACCUGAGUGGUCUGUAUUUAAUUUAUAGUUCCCAAUUCGGUAGC
GGUUGUAUGCAGCAAAACCUGAAUCGUUGCCUACACGCUGCGAAGCUCCUAAUUUGUAAUAAGAAAGC
GUUCGUGAUGUAGCCACCGUAAUCUCUUUUGGCAGGUCCUUUAUGUCACAGCGCCCUAGGGAGUGUCC

>Orf 7 Subgenomic

AAAGCAAAAUGUGUGCUAAUGCAAGUUAGUGCAAAUUUGUUGUCAGCAAGAGGAUGAAAUGGUGAAU
UGCCCUCGUAGGUCCCUGACGGGCAAGGUUCUUUUAGUAGUACAGUGGUACCUCUAACACACUCCUGA
UAGUGAUAUAGCUCGCUGGAAGCAAGUGCAAUCAAUGUCAAGAAGAGAAUAAUUUUCAUGUUCGUUU
UAUGGAUAAUCUAACUCCAUAGGUUCUUCAUCAUCUAACUCAGAGUAUUUCUUCUUAGUUAGAGGCUU
AAAUAAUUGUCUCACUAUUGAACUUAUUAGUAUGUCAAGGUUCCAAAUGGCAACCCUGAAAGUCUUCA
UAAUGAUAAUCAAUAUCUCUGCUAUUGUAACCUGGAAGUCAACAAGAUGAAACAUCUGUUGUCACUUA

>Orf 8 subgenomic

AUUGUACAAGGCGAGCUGACUUUCUAGAUCCAAUUUUUAUAAACCAGUCCGAAUAGUAAUGUAUUGGA
CAUGGGUCUUCAAUUUGGUAGGGUUGAUUUUCACAACACUCUUGUAUGCUGCAUUCUUUAUGAAUGC
AGUAUACUGAUGUUAAGAGUCCAAAAACAAUGAGAAGUUUCAUGUUCGUUUAGACUUUGUUACAGGG
UUCUUCUAUAUCCUGGAGUUCAAGUGAAAACCAAAAUAUAAUAAGCAUUAUUAGAACAAGGAAUAGCA
GAAAGGCUAAAAAGCACAAAUAGAAGUCAAUUAAAGUGAGCUCAUUCAUUCUGUCUUUCUCUUAAUGG
UGAAGCAAAGUAUUAUAAAUACUAGAGCAGCAACAAUGAGAAAAAGUGGUGAAUAGAGCUCUUGGUGA

>N subgenomic

AGUAUUAUUGGGUAAACCUUGGGGACGGCGUUGCUUUGGCCGUGCACCACUGCGUCCUCCAUCCUGAU
UAUUGUCAGUUGAAUCUGAGGGUCCACCAAAUGUAAUGCGGGGGGCACUACGCUGGUUUUGGGGUCC

AUUAUCAGACAUUUUAGUUUGUUCGUUUAGAUGAAAUCUAGAACAACACGCACGUCAUGAUGCUCAAC AAAGUCAUAAUCGUACGAGCAGCGCACAAUUAGACUACCUACUGGUGGUGCUUGACAGUUUAUCUCAA GGGGUUCACAAGAGAUAGUGUAAUUGCCAAACAUUUCAUAAUGAAUCGGAAUUCUUUUGCCAUAAUCU CCCUCGCACAAUUGUACAAGGCGAGCUGACUUUCUAGAUCCAAUUUUUAUAAACCAGUCCGAAUAGUAA

**Sequences used for secondary structure prediction of SARS-CoV-2 subgenomic RNAs by using Vienna RNA webserver:**

**Specific length of aligned RNA sequence was used for secondary structure prediction**

>S Subgenomic

AACUGAGGAUCUGAAAACUUUGUCAGGGUAAUAAACACCACGUGUGAAAGAAUUAGUGUAUGCAGGGG GUAAUUGAGUUCUGGUUGUAAGAUUAACACACUGACUAGAGACUAGUGGCAAUAAAACAAGAAAAACA AACAUUGUUCGUUUAGUUGUUAACAAGAACAUCACUAGAAAUAACAACUCUGUUGUUUUCUCUAAUUA UAAGUCUACCUUUACUAAGAAGAGAUAAAAUCAUAUCAUUGAUUUGACCUUCUUUUAAAGACAUAACA GCAGUACCCCU

>Orf3a Subgenomic

AGCGCGAACAAAAUCUGAAGGAGUAGCAUCCUUGAUUUCACCUUGCUUCAAAGUUACAGUUCCAAUUG UGAAGAUUCUCAUAAACAAAUCCAUAAGUUCGUUUAUGUGUAAUGUAAUUUGACUCCUUUGAGCACUG GCUCAGAGUCGUCUUCAUCAAAUUUGCAGCAGGAUCCACAAGAACAACAGCCCUUGAGACAACUACAGC AACUGGUCAUACAGCAAAGCAUAAUUGUCACCAUUACUAUGGCAAUCAAGCCAGCUAUAAAACCUAGCC AAAUGUACCAU

>E subgenomic

AGAAUACCACGAAAGCAAGAAAAAGAAGUACGCUAUUAACUAUUAACGUACCUGUCUCUUCCGAAACGA AUGAGUACAUAAGUUCGUACUCAUCAGCUUGUGCUUACAAAGGCACGCUAGUAGUCGUCGUCGGUUCA UCAUAAAUUGGUUCCAUUACUGGAUUAACAACUCCGGAUGAACCGUCGAUUGUGUGAAUUUGGACAUG UUCUUCAGGCUCAUCAACAAUUUUAUUGUAGAUGAAGAAGGUAACAUGUUCAACACCAGUGUCUGUAC UCAAUUGAGU

>M subgenomic

UAGGAAACCUAUUACUAGGUUCCAUUGUUCAAGGAGCUUUUUAAGCUCUUCAACGGUAAUAGUACCGU UGGAAUCUGCCAUGGCUAAAAUUAAAGUUCCAAACAGAAAAACUAAUAUAAUAUUUAGUUCGUUUAGA CCAGAAGAUCAGGAACUCUAGAAGAAUUCAGAUUUUUAACACGAGAGUAAACGUAAAAAGAAGGUUUU ACAAGACUCACGUUAACAAUAUUGCAGCAGUACGCACACAAUCGAAGCGCAGUAAGGAUGGCUAGUGUA ACUAGCAAGA

> Orf6a Subgenomic

AGGUUUAUGAUGUAAUCAAGAUUCCAAAUGGAAACUUUAAAAGUCCUCAUAAUAAUUAGUAAUAUCUC UGCUAUAGUAACCUGAAAGUCAACGAGAUGAAACAUCUGUUGUCACUUACUGUACAAGCAAAGCAAUA UUGUCACUGCUACUGGAAUGGUCUGUGUUUAAUUUAUAGUUGCCAAUCCUGUAGCGACUGUAUGCAG CAAAACCUGAGUCACCUGCUACACGCUGCGAAGCUCCCAAUUUGUAAUAAGAAAGCGUUCGUGAUGUAG CAACAGUGAUUUC

>Orf7a subgenomic

CGUAUGUUCCAGAAGAGCAAGGUUCUUUUAAAAGUACUGUUGUACCUCUAACACACUCUUGGUAGUGA
UAAAGCUCACAAGUAGCGAGUGUUAUCAGUGCCAAGAAAAGAAUAAUUUUCAUGUUCGUUUAAUCAAU
CUCCAUUGGUUGCUCUUCAUCUAAUUGAGAAUAUUUAUUCUCAGUUAGUGACUUAGAUAAAUUUUUA
AUUAUGAGGUUUAUGAUGUAAUCAAGAUUCCAAAUGGAAACUUUAAAAGUCCUCAUAAUAAUUAGUAA
UAUCUCUGCUAUA

>Orf7b subgenomic

UGAUCUUGCAGUUCAAGUGAGAACCAAAAGAUAAUAAGCAUAAUUAAAACAAGGAAUAGCAGAAAGGC
UAAAAAGCACAAAUAGAAGUCAAUUAAUGAAAGUUCAAUCAUUCUGUCUUUCUUUUGAGUGUGAAGCA
AAGUGUUAUAAACACUAUUGCCGCAACAAUAAGAAAAAUUGGAGAGUAAAGUUCUUGAACUUCCUCUU
GUCUGAUGAACAGUUUAGGUGAAACUGAUCUGGCACGUAACUGAUAGACGUGUUUUACGCCGUCAGGA
CAAGCAAAAGCAA

> Orf 8b subgenomic

ACACGGGUCAUCAACUACAUAUGGUUGAUGUUGAGUACAUGACUGUAAACUACAUUCUUGGUGAAAUG
CAGCUACAGUUGUGAUGAUUCCUAAGAAAACAAGAAAUUUCAUGUUCGUUUAGGCGUGACAAGUUUCA
UUAUGAUCUUGCAGUUCAAGUGAGAACCAAAAGAUAAUAAGCAUAAUUAAAACAAGGAAUAGCAGAAA
GGCUAAAAAGCACAAAUAGAAGUCAAUUAAUGAAAGUUCAAUCAUUCUGUCUUUCUUUUGAGUGUGAA
GCAAAGUGUUAU

>N subgenomic

GUUACUGCCAGUUGAAUCUGAGGGUCCACCAAACGUAAUGCGGGGUGCAUUUCGCUGAUUUUGGGGU
CCAUUAUCAGACAUUUUAGUUUGUUCGUUUAGAUGAAAUCUAAAACAACACGAACGUCAUGAUACUCU
AAAAAGUCUUCAUAGAACGAACAACGCACUACAAGACUACCCAAUUUAGGUUCCUGGCAAUUAAUUGUA
AAAGGUAAACAGGAAACUGUAUAAUUACCGAUAUCGAUGUACUGAAUGGGUGAUUUAGAACCAGCCUC
AUCCACGCACAA

**Supplementary Figures: Secondary structure of negative sense subgenomic RNAs (sequences are provided in page number 3-4)**

**Figure 1: E subgenomic**

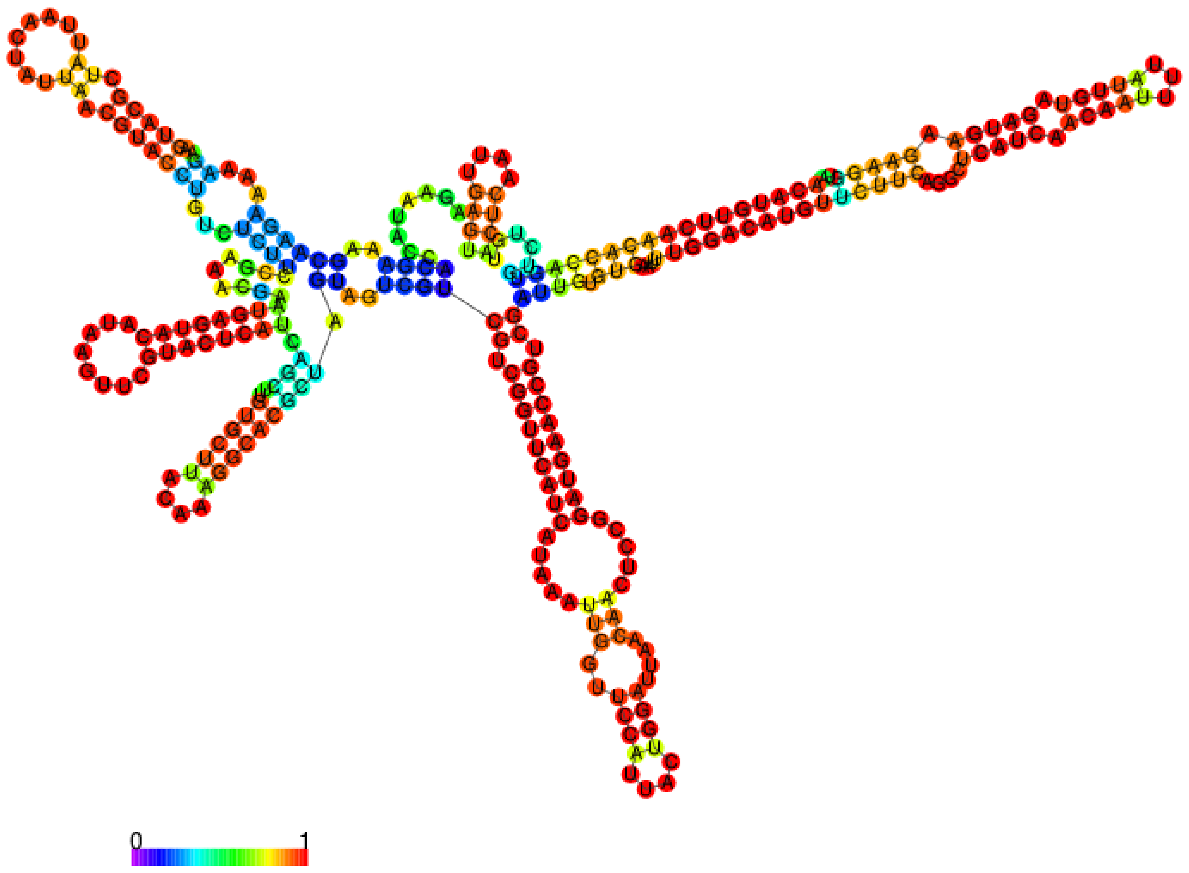


**Figure 2: M subgenomic**

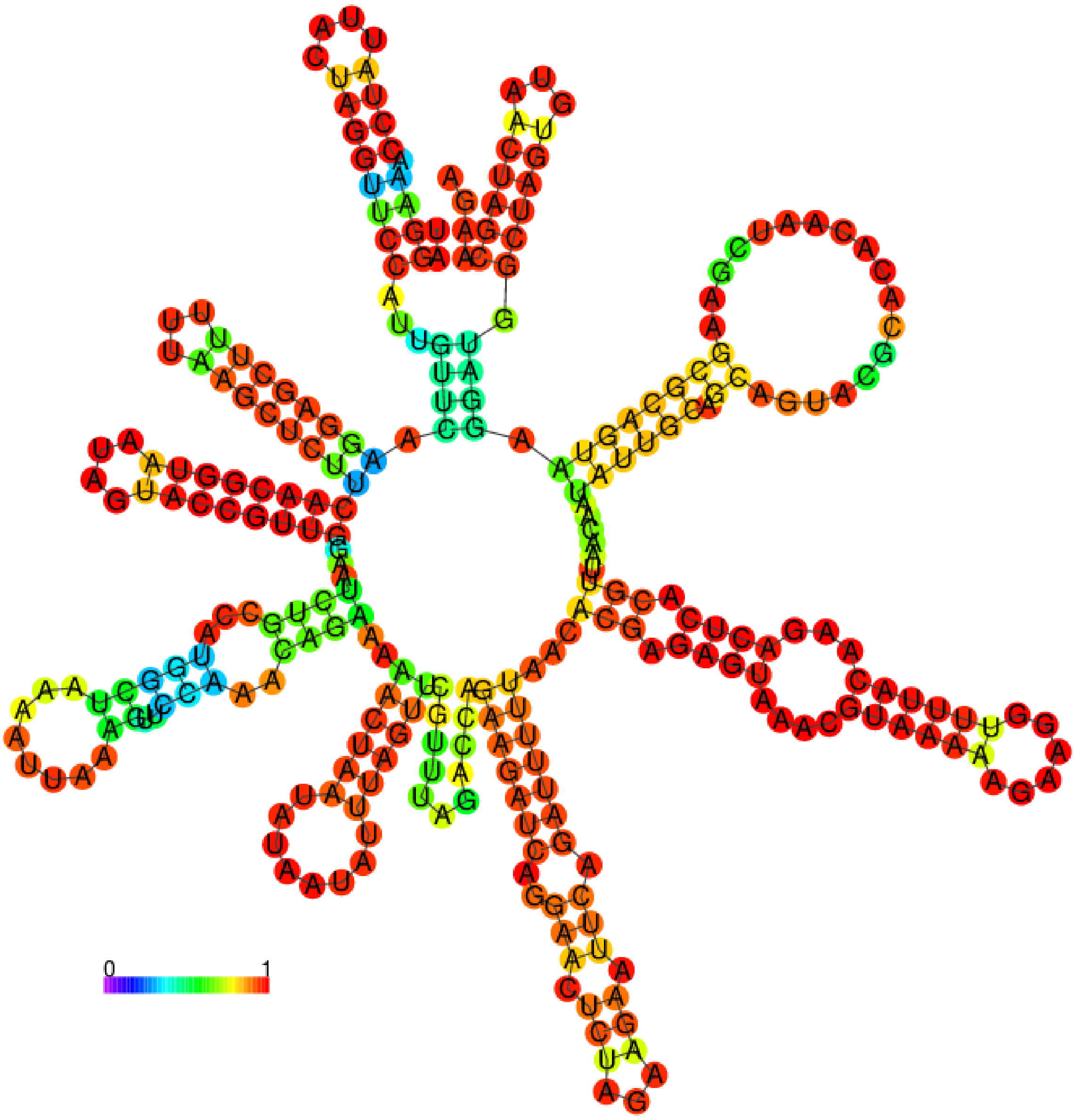


**Figure 3: N subgenomic**

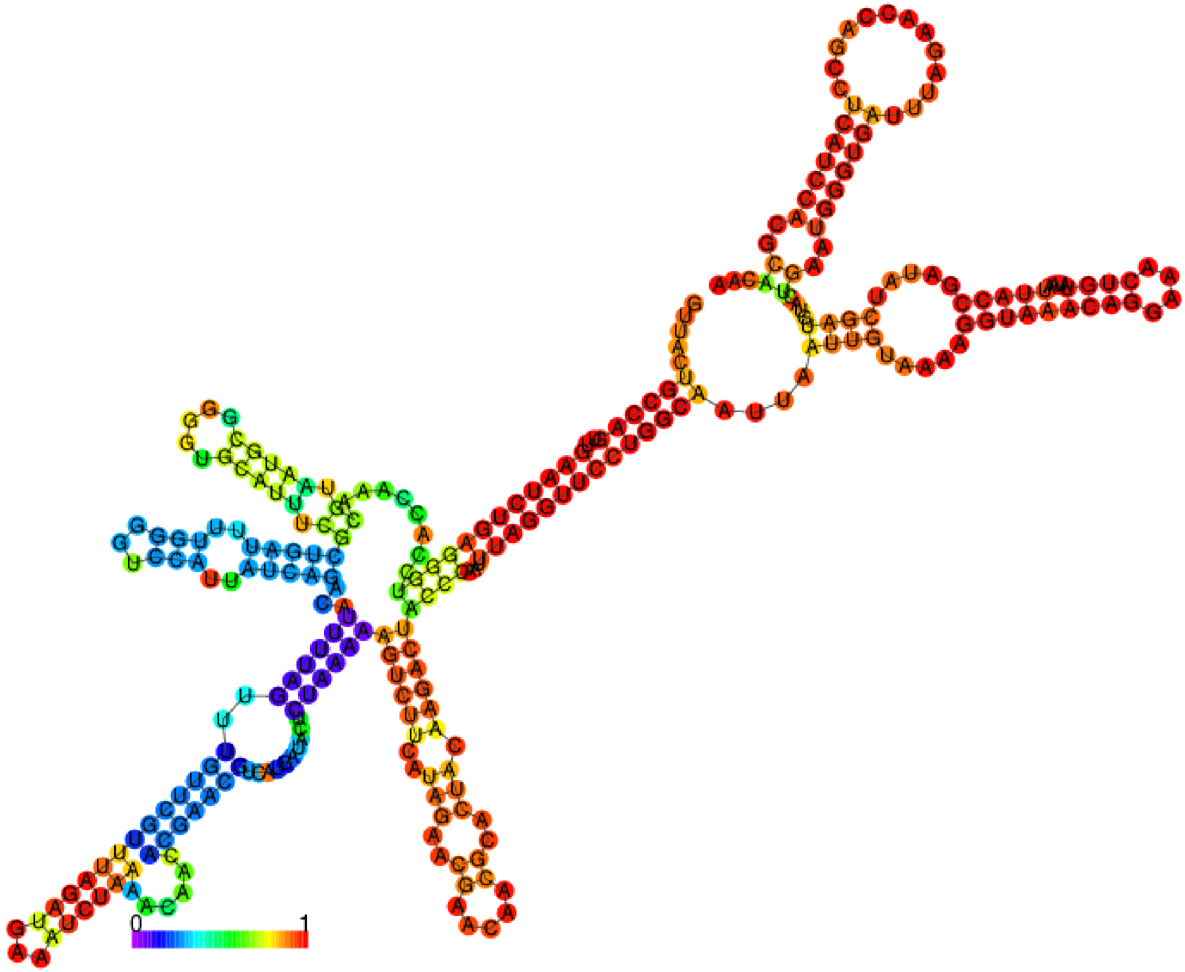


**Figure 4: Orf3a Subgenomic**

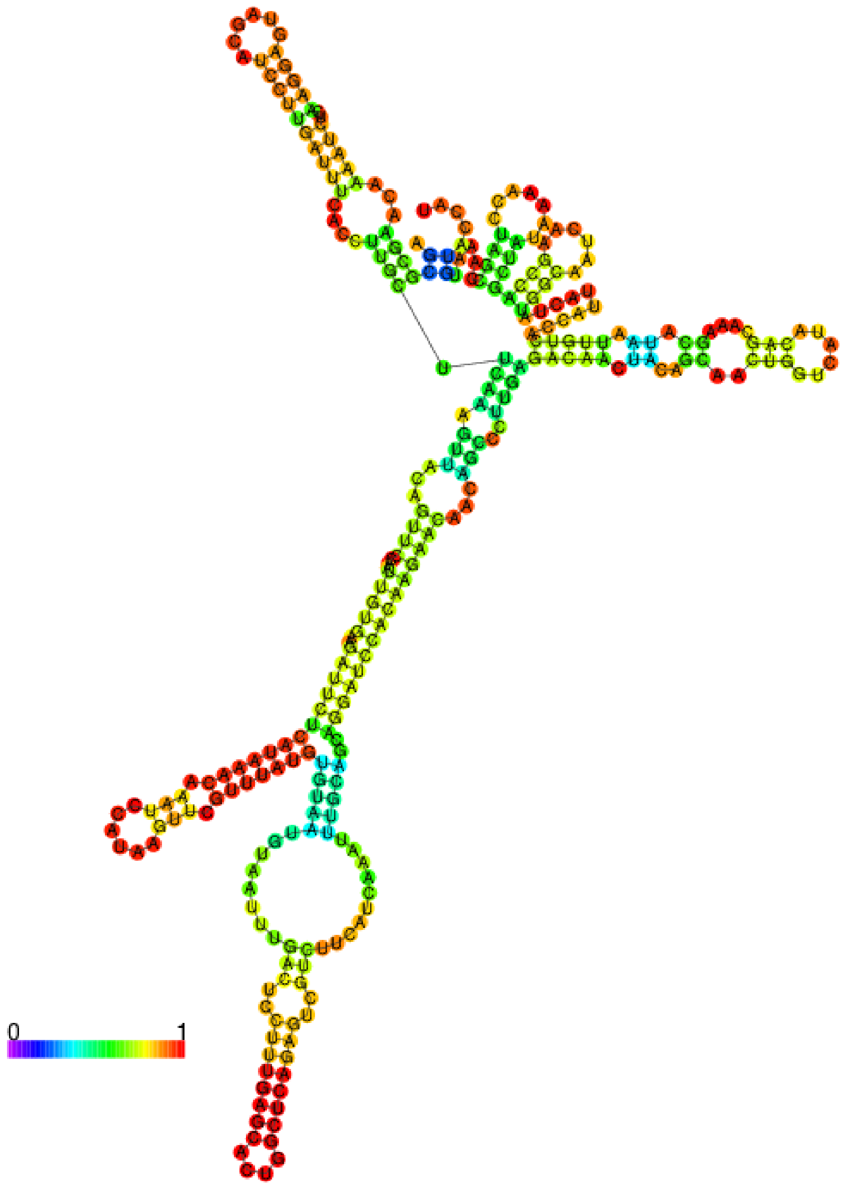


**Figure 5: Orf 6 Subgenomic**

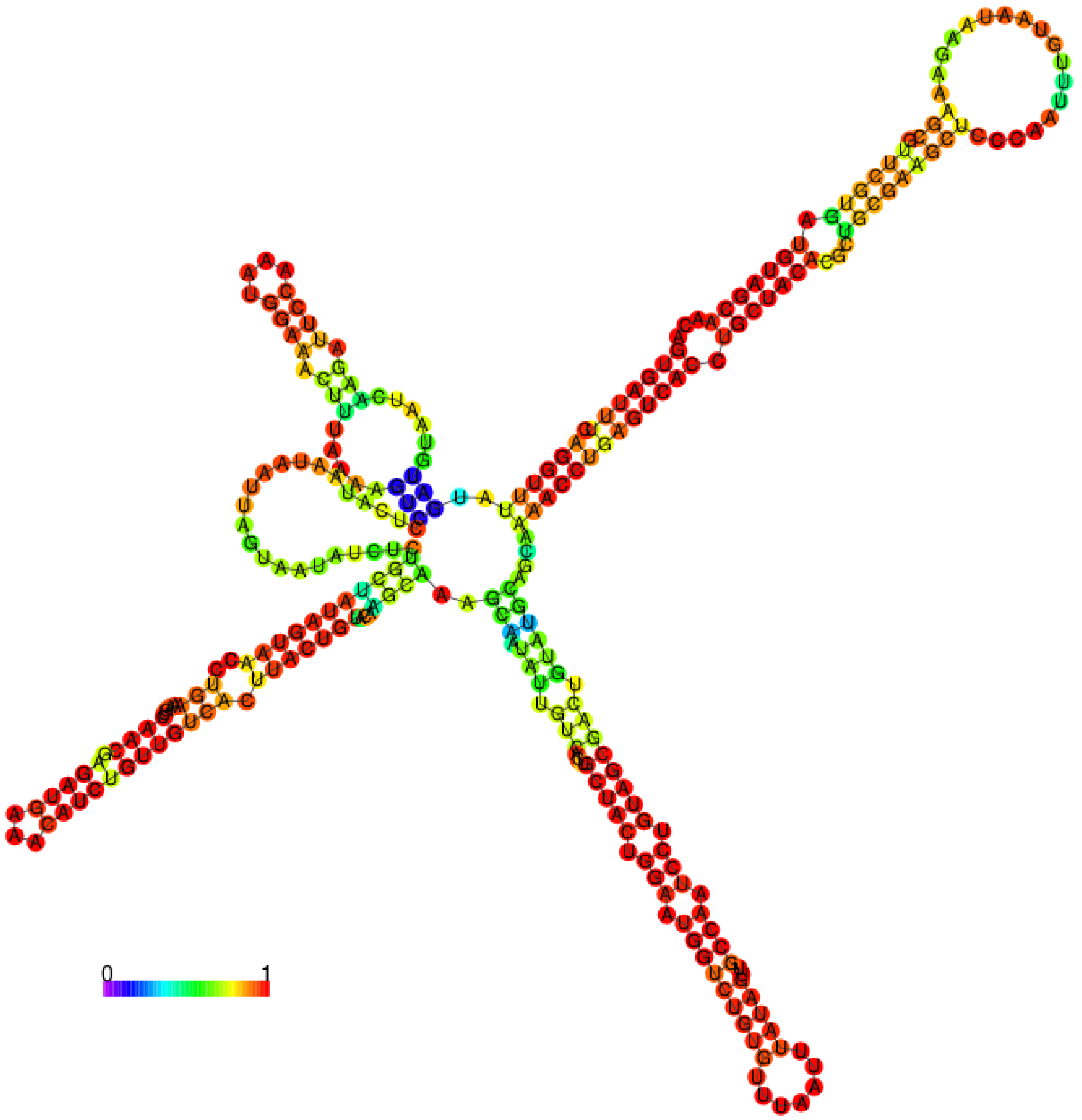


**Figure 6: Orf 7a subgenomic**

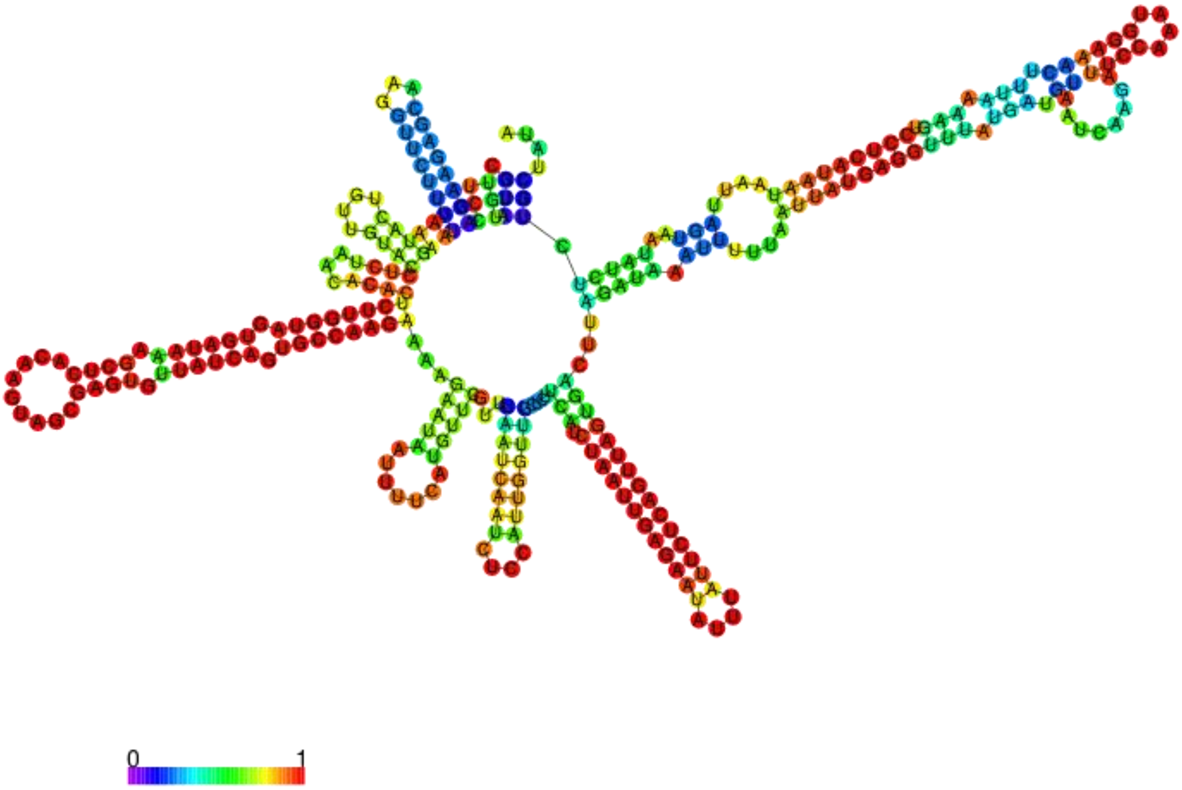


**Figure 7: Orf 7b subgenomic**

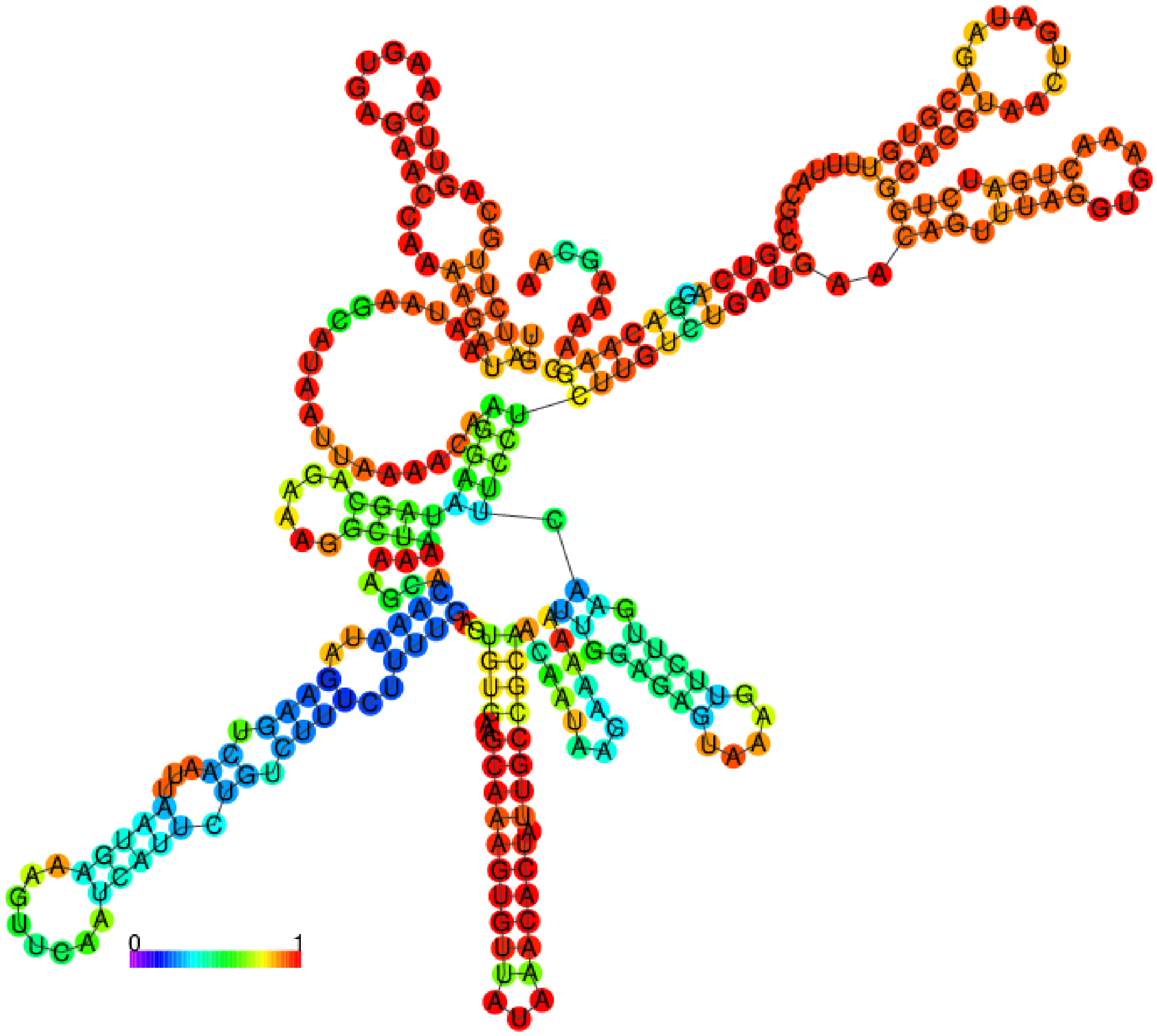


**Figure 8: Orf 8 subgenomic**

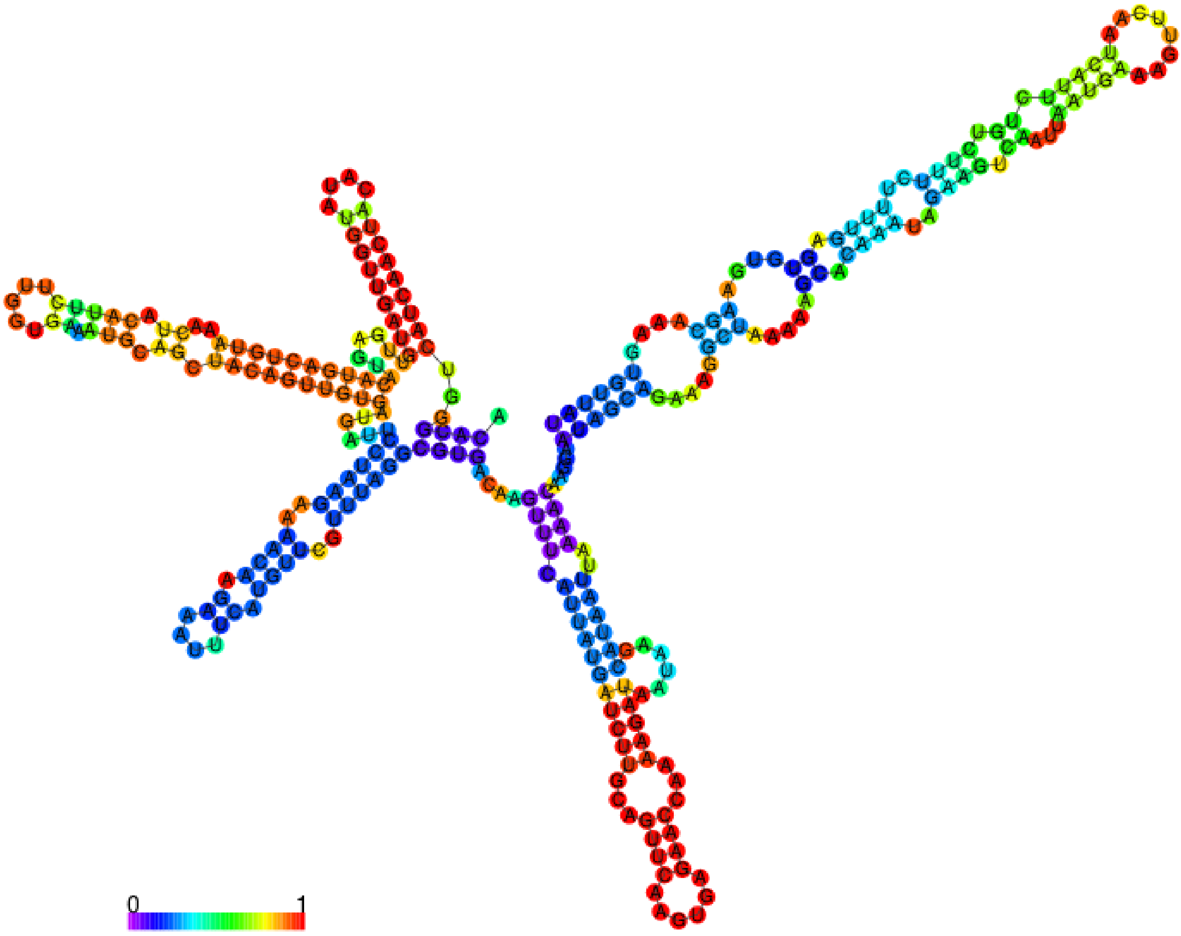


**Figure 9: Orf S subgenomic**

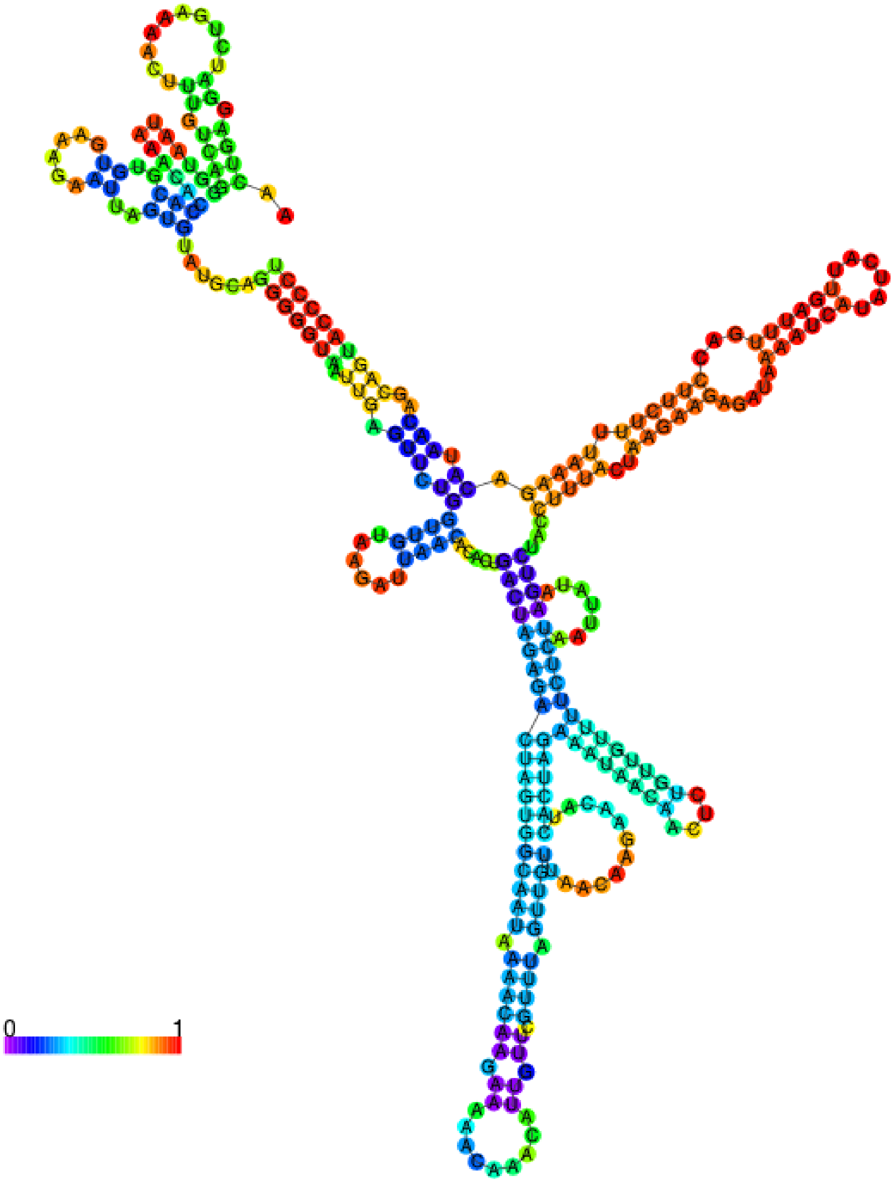
0
1
